# Modulation of Arterial Intima Stiffness by Disturbed Blood Flow

**DOI:** 10.1101/2023.09.07.556773

**Authors:** Briana C. Bywaters, Andreea Trache, Gonzalo M. Rivera

## Abstract

**Background:** The intima, comprising the endothelium and the subendothelial matrix, plays a crucial role in the development of atherosclerotic plaques, especially in bifurcations and curved segments of arteries. The mechanical stress arising from disturbed blood flow (d-flow) and the stiffening of the arterial wall contributes to endothelial dysfunction. However, the specific impacts of these physical forces on the mechanical environment of the intima remain undetermined. To address this gap in knowledge, we investigated whether inhibiting collagen crosslinking could ameliorate the detrimental effects of persistent d-flow on the mechanical properties of the intima.

**Methods:** To explore this hypothesis, we performed partial ligation (PCL) of the left carotid artery (LCA) in male and female C57BL/6J mice, inducing d-flow. The right carotid artery (RCA) served as an internal control. Carotids were collected two days and two weeks after PCL to study acute and chronic effects of d-flow on the mechanical phenotype of the intima. To decouple the chronic effects of d-flow from the ensuing arterial wall stiffening, we used subcutaneous implants delivering either phosphate-buffered saline (Saline) or 150 mg/kg/day of β-aminopropionitrile (BAPN), an inhibitor of elastin and collagen crosslinking lysyl oxidase (LOX) and LOX-like (LOXL) enzymes. Atomic force microscopy (AFM) measurements allowed us to determine stiffness of the endothelium and the denuded subendothelial matrix in *en face* carotid preparations. In addition, we determined the stiffness of human aortic endothelial cells (HAEC) cultured on soft and stiff hydrogels.

**Results:** Acute exposure to d-flow caused a slight decrease in endothelial stiffness in male mice but had no effect on the stiffness of the subendothelial matrix in either sex. Regardless of sex, the intact endothelium was softer than the subendothelial matrix. In contrast, exposure to chronic d-flow led to a substantial increase in the endothelial and subendothelial stiffness in both sexes. The effects of chronic d-flow were largely prevented by concurrent BAPN administration. Notably, the subendothelial matrix of ligated, BAPN-treated arteries was softer than that of unligated, saline-treated counterparts. Furthermore, HAEC displayed reduced stiffness when cultured on soft vs. stiff hydrogels.

**Conclusions:** Exposure to chronic d-flow results in marked stiffening of arterial intima, which can be effectively prevented by pharmacological inhibition of LOX/LOXL enzymes.

**Highlights:**

- Acute exposure to d-flow slightly softens the endothelium in males.
- Chronic exposure to d-flow causes stiffening of the arterial intima.
- Inhibition of LOX/LOXL enzymes prevents intimal stiffening arising from chronic d-flow.

## Introduction

Atherosclerotic lesions primarily develop in areas where arteries branch out or have curved segments,^1^ despite the influence of systemic risk factors. The mechanical forces exerted by local blood flow patterns play a crucial role in determining the endothelial phenotype and influencing the progression of atherosclerosis.^2, 3^ In straight arterial regions, the high-magnitude and unidirectional shear stress resulting from steady blood flow promotes endothelial health and is considered protective against atherosclerosis. On the other hand, in bifurcations and curved arterial segments, the low-magnitude and oscillatory shear stress caused by disturbed blood flow (d-flow) leads to endothelial dysfunction and is deemed prone to atherosclerosis.

The mechanical properties of the arterial wall also play a significant role in the development and progression of atherosclerosis.^4^ Factors such as aging,^5^ metabolic imbalance,^6^ and hypertension,^7^ contribute to arterial stiffening by causing changes in the extracellular matrix and vascular cells. Arterial stiffening is also observed in bifurcations and curved segments, which aligns with the localized formation of atherosclerotic lesions due to d-flow.^8-10^ However, the specific impact and interplay between chronic exposure to d-flow and arterial wall stiffening on the intimal phenotype, and specifically its biomechanical properties, remain poorly understood.

The mouse model of partial carotid ligation (PCL) has been valuable in uncovering the mechanisms involved in the development of atherosclerosis.^11-13^ By surgically inducing d-flow in a previously healthy and straight arterial segment, the development of atherosclerotic lesions can be rapidly stimulated in hyperlipidemic mice.^11, 14, 15^ In the presence of physiological levels of cholesterol and triglycerides, however, prolonged exposure (∼two weeks) to PCL-induced d-flow leads to arterial wall remodeling and stiffening, similar to the effects of aging.^16^ This model offers, therefore, a unique opportunity to study how mechanical forces contribute to abnormal molecular patterns and cellular phenotypes that drive the progression of atherosclerosis.^13^ This research aimed to ascertain the specific contribution of d-flow and arterial wall stiffening to the mechanical phenotype of components of the arterial intima, i.e., the intact endothelium and the denuded subendothelial matrix. The hypothesis tested was that inhibition of arterial wall stiffening neutralizes the deleterious effects of chronic d-flow on the mechanical properties of the arterial intima. To test this hypothesis, we combined d-flow induced by PCL with a pharmacological approach to reduce arterial wall stiffness. Our findings indicate that reducing arterial wall stiffness can reverse the detrimental effects of prolonged exposure to d-flow on the biomechanical properties of the arterial intima.

## Methods

### Animals and experimental design

We performed partial ligation of the left carotid artery (PCL)^11, 12^ in C57BL/6J mice to induce d-flow (Suppl. Fig. 1A). To verify arterial wall integrity and the subsequent removal of the endothelium (Suppl. Fig. 1C, D) for determination of endothelial and subendothelial matrix stiffness, respectively, we used the fluorescence reporter mT/mG mouse^17^ expressing tamoxifen-inducible Cre recombinase (*Cdh5*-Cre^ERT2^) in the endothelium.^18^ In these mice, expression of membrane-tagged tdTomato is constitutive in all tissues. Upon treatment with tamoxifen, Cre-mediated recombination occurs only in endothelial cells which results in the expression of membrane-tagged enhanced green fluorescent protein (mEGFP). To determine the acute and chronic effects of d-flow on the mechanical phenotype of the endothelium and subendothelial matrix, animals were sacrificed, and carotid arteries collected 2 days or 2 weeks after PCL (Suppl. Fig. 1B). To decouple the effects of chronic (2 weeks) d-flow from the ensuing arterial wall stiffening, animals were affixed with subcutaneous implants delivering either phosphate buffered saline (Saline) or 150 mg/kg/day of β-aminopropionitrile (BAPN), an inhibitor of LOX/LOXL enzymes (Suppl. Fig. 1B). Mice were maintained in the Comparative Medicine Program Laboratory Animal Resources and Research facility at Texas A&M University. Animal handling and experimental procedures were performed according to a protocol (AUP 2019-0184) approved by the Institutional Animal Care and Use Committee of Texas A&M University. Mice were maintained on a 12-h light/dark cycle and fed regular chow ad-libitum. Both male and female mice were used in the study.

### Surgical procedures

At approximate 8 weeks of age, mice were anesthetized with isoflurane and hair was chemically removed (Nair, Ewing, NJ) from the chin to the bottom of the sternum. The exposed skin was swabbed alternately with iodine and isopropyl alcohol prior to incision. Sutures (0-6 silk) were used to occlude the external carotid, internal carotid, and occipital branches of the left carotid artery (LCA), leaving the superior thyroid artery unobstructed (Suppl. Fig. 1A). The right carotid artery (RCA) was left intact and served as an internal control. The incision was closed using tissue adhesive (Vetbond, 3M, Saint Paul, MN). Soon after surgery, all mice received a subcutaneous injection of buprenorphine (0.1 mg/kg) to minimize pain and discomfort. Following PCL, mice used for the two-week experiments remained anesthetized and were placed in a prone position. Hair was chemically removed from between the shoulder blades and the skin was swabbed with iodine and isopropyl alcohol. A small incision was made to allow the subcutaneous insertion of an osmotic minipump (ALZET model 1002, capacity: 100 µL, flow rate: 0.25 µL/h) loaded with either Saline or BAPN, which was previously dissolved in phosphate buffered saline at 37°C. Following the subcutaneous insertion of the osmotic minipump, the incision was closed using 7mm wound clips (EZ Clip, Stoelting, Chicago, IL).

### Doppler ultrasound

We used ultrasound to confirm successful PCL one day after the surgical procedure. Mice were anesthetized using isoflurane and subjected to Doppler ultrasound using a Vevo 3100 system (FUJIFILM VisualSonics, Inc., Bothell, WA) in B-mode. Both flow reversal and decreased flow velocity were recorded in the ligated vessels.^11, 12^ The presence of normal flow was confirmed in the intact vessels (Suppl. Fig. 1A, bottom panels).

### Tissue collection

Two days or two weeks following PCL, mice were euthanized using CO2 asphyxiation followed by cervical dislocation. The vascular tree was perfused through the apex of the heart with ice-cold Dulbecco’s phosphate buffered saline (DPBS, Gibco, Waltham, MA) prior to excision of carotid arteries. Excised vessels were immediately opened *en face* using vannas scissors (Fine Science Tools, Vancouver BC, Canada), affixed to 60 mm plastic dishes (BD Falcon, Franklin Lakes, NJ), and submerged in room temperature DPBS.

### Culture of endothelial cells

Human aortic endothelial cells (ATCC, Manassas, VA) were cultured in EGM-2 culture media (Lonza Walkersville, Basel, Switzerland) on 0.1% gelatin-coated plastic dishes until ∼80% confluence. Cells were then seeded (passage 5) on soft (4 kPa) or stiff (50 kPa) polyacrylamide hydrogels coated with 0.2% collagen I (Matrigen, San Diego, CA) at a density of 40x10^4^/18mm^2^ and cultured for 24 hours. Experiments were performed in triplicates.

### Atomic force microscopy

To determine the stiffness of the intact endothelium and the subendothelial matrix, we performed atomic force microscopy (AFM) as previously described.^19^ The AFM probes used were unsharpened silicon nitride cantilevers with a spring constant of 12.2 ± 0.4 pN/nm (MLCT Bio, Bruker Nano-Surfaces, Billerica, MA). The AFM was operated in force mode, setting the cantilever to touch and retract from the tissue surface at 0.8 µm/s in the z-axis. Stiffness at the point of contact was evaluated as Young’s modulus of elasticity (E), by fitting the approach curve between the initial point of tissue contact and point of maximum probe displacement with Sneddon’s modified Hertz model.^19^ On each vessel, 4-8 separate sites were measured and approximately 60 force curves were acquired at each site. Care was taken to avoid obtaining measurements at the edges of the tissue due to eventual damage during handling and immobilization in the dish. After measurement of the intact endothelial surface was completed, the endothelial layer was removed using 10 swipes with a cotton swab (Suppl. Fig. 1C, D). DPBS was replaced and the exposed subendothelial matrix of the vessels was subjected to AFM measurements using the same approach described for the intact endothelium.

In the case of HAEC cultured on hydrogels, measurements of cells stiffness and integrin α5β1 adhesion force to fibronectin were conducted using AFM tips coated with fibronectin.^20, 21^ Individual cells were measured midway between the nucleus and the edge of the cytoplasm.^22^ Approximately 10 cells were measured, for a total of 1330-1344 force curves per condition.

### AFM data processing and statistical analysis

Kernel density plots of the distribution of cell stiffness at the point of contact for both *ex vivo* vessels and cultured cells, as well as kernel density plots of the distribution of integrin adhesion force measurements for cultured cells, were generated using NForceR software.^23^ These probability density plots were then analyzed using Fityk^24^ to generate 95% confidence intervals of peak values based on the Lorentzian model. Peaks with non-overlapping confidence (CI) intervals were deemed statistically different (*p*<0.05). In addition, we determined half-width half-max (HWHM) values for each fitted distribution as an estimate of stiffness heterogeneity in a sample. The HWHM parameter describes the full width of a distribution at half of its maximum high, i.e., it is the distance between the two points on the x-axis where the probability density function of the distribution drops to half of its maximum value.

To compare variability between individual vessels, mean (average?) stiffness values were calculated across the 4-8 sites/vessel for each experimental condition.^25^ Statistical significance (*p*<0.05) between vessels (RCA, LCA) harvested from animals that underwent two days of PCL (Suppl. Fig. 2) was determined using Student’s t-test. Statistical significance (*p*<0.05) between vessels (RCA+saline, LCA+saline, LCA+BAPN) harvested from animals that underwent two weeks of PCL (Suppl. Fig. 4) was determined using ANOVA followed by Tukey post-hoc testing with Bonferroni correction.

## Results

### Effects of acute exposure to d-flow on the mechanical properties of the carotid intima

To determine how acute d-flow affects the mechanical properties of the carotid intima, we harvested carotid arteries from male and female mice (n=6/sex) two days following PCL. We used AFM to measure the stiffness of the intact endothelium and subendothelial matrix in both the intact RCA and ligated LCA. In males, acute exposure to d-flow slightly decreased (p<0.05) the endothelial stiffness (Fig. 1A) but had no effect on the stiffness of the subendothelial matrix (Fig. 1B). Acute exposure to d-flow in females, on the other hand, did not affect (*p*>0.05) the endothelial (Fig. 1C) or subendothelial stiffness (Fig. 1D). No statistical differences were detected in either males or females when the mean stiffness of unligated RCAs was compared to that of ligated LCAs (Suppl. Fig. 2). Regardless of sex, the intact endothelium was softer (*p*<0.05) than the subendothelial matrix.

**Figure 1.**
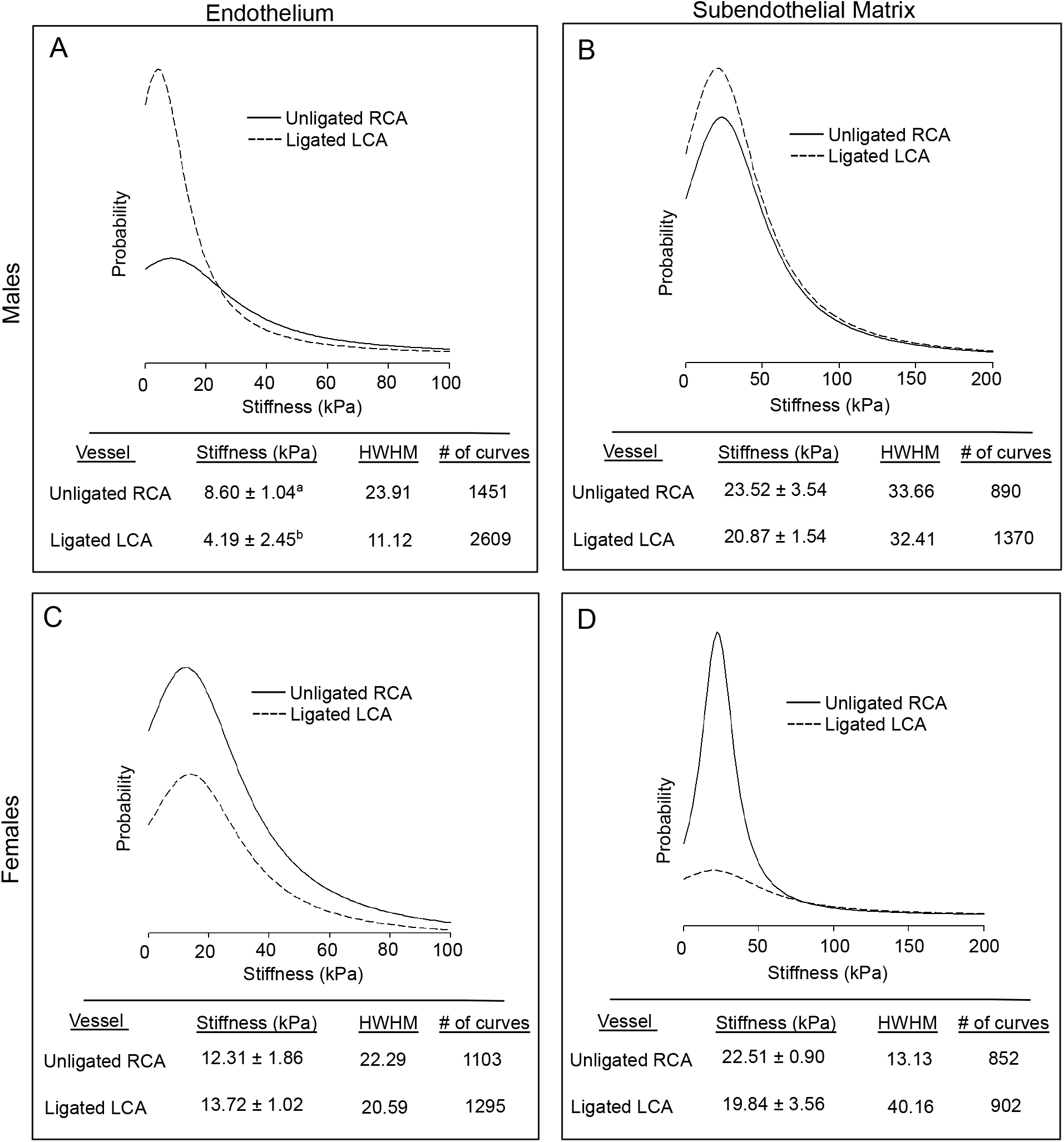
Effects of acute exposure to d-flow on the stiffness of the carotid intima. The stiffness of the endothelium (A,C) and subendothelial matrix (B,D) was measured using AFM in male and female mice 2 days after PCL. In each panel, force curves fitted with a Lorentzian model are shown (top left). Fitted force curves were generated by combining data for unligated RCAs (solid line) or ligated LCAs (dashed line) from all animals (n=6/sex). The table summarizes stiffness values (E peak ± 95% confidence interval, kPa) and number of force curves (bottom). HWHM, half-width half-max. a,b, p<0.05.

To assess heterogeneity across the vessel, the half-width half-max (HWHM) value of the Lorentzian curve fit for each vessel type (unligated RCA, ligated LCA) was compared. In males, there was a ligation-induced decrease in the endothelial HWHM (Fig. 1A, table) but no changes were observed for this index in the subendothelial matrix (Fig. 1B, table). In females, the endothelial HWHM remained unchanged (Fig. 1C, table) but, in response to ligation, this index increased ∼3 fold in the subendothelial matrix (Fig. 1D, table). Suppl. Fig. 3 shows the Lorentzian curve fittings (solid line) used to determine peak values and 95% CIs of stiffness measurements (dotted line, n=852 to 2,609/per condition) from arteries harvested 2 days after PCL.

### Effects of chronic exposure to d-flow on the mechanical properties of the carotid intima

How chronic (2 weeks) d-flow affects the mechanical properties of the carotid intima in male and female mice treated with saline or BAPN (n=6/sex/condition) was also determined. Treatment with BAPN was implemented to decouple the effects of chronic d-flow from the ensuing stiffening of the arterial wall. In saline-treated mice, regardless of sex, chronic d-flow induced a ∼8-fold increase (p<0.05) in the stiffness of the endothelium (Fig. 2A, C). Such response was reduced, to a large extent, by the administration of BAPN. Although BAPN effectively prevented ligation-induced endothelial stiffening, the endothelial stiffness of ligated LCA+BAPN carotids was ∼2 fold higher than that of unligated RCA+saline counterparts. The increase in the endothelial HWHM value observed in response to chronic d-flow (Fig. 2A, C, table) was largely offset by BAPN treatment in males but not females.

**Figure 2.**
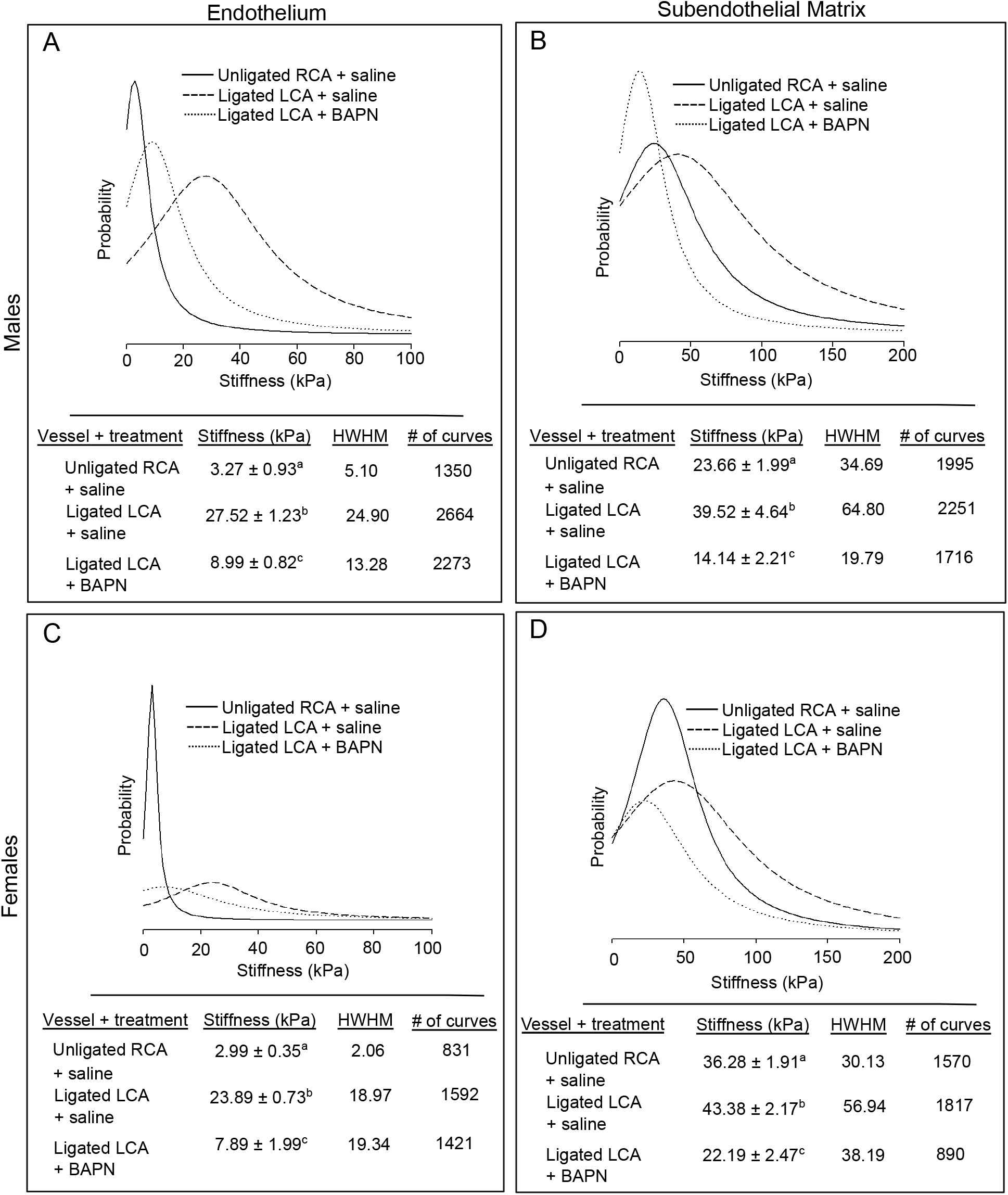
Effects of chronic exposure to d-flow on the stiffness of the carotid intima. The stiffness of the endothelium (A,C) and subendothelial matrix (B,D) was measured using AFM in male and female mice 2 weeks after PCL and treatment with either sterile phosphate-buffered saline (Saline) or 150 mg/kg/day of β-aminopropionitrile (BAPN), an inhibitor of the collagen crosslinking enzymes of the lysyl oxidase family. Within each panel, force curves fitted with a Lorentzian model are shown (top left). Fitted force curves were generated by combining data for unligated RCA treated with saline (solid line), ligated LCA treated with saline (dashed line), and ligated LCA treated with BAPN (dotted line) from all animals (n=6/sex/group). The table summarizes stiffness values (E peak ± 95% confidence interval, kPa) and number of force curves (bottom). HWHM, half-width half-max. a,b,c, p<0.05.

Chronic exposure to d-flow stiffened the subendothelial matrix in both sexes, an effect that was fully counteracted (p<0.05) by concurrent BAPN treatment (Fig. 3B and D). Noticeably, the subendothelial matrix of LCA+BAPN arteries was softer than that of RCA+saline carotids. In addition, chronic d-flow almost doubled the subendothelial matrix HWHM value (Fig. 3B and D, table). Such effect, however, was prevented by BAPN treatment.

Comparison among groups of vessels showed that the increase in the mean stiffening of the endothelial and the subendothelial matrix induced by chronic d-flow was blocked by BAPN treatment (Suppl. Fig. 4). Suppl. Fig. 5 shows the Lorentzian curve fittings (solid line) used to determine peak values and 95% confidence intervals of stiffness measurements (dotted line, n= 831 to 2,664/per condition) from arteries harvested 2 weeks after PCL. Noticeably, the AFM cantilever used in these studies was not soft enough to capture force curve profiles in samples of unligated RCA+BAPN carotids that, unexpectedly, presented a very soft nature.

### The mechanical properties of endothelial cells depend on substrate stiffness

To determine the effect of substrate stiffness on human aortic endothelial cells (HAEC) *in vitro*, cells were seeded on collagen-coated polyacrylamide hydrogels. Soft (4 kPa) and stiff (50 kPa) hydrogels were used to mimic, respectively, the mechanical properties of the healthy and diseased arterial wall.

Twenty-four hours after seeding, stiffness and α5β1 integrin-fibronectin adhesion force at the point of contact was measured in live cells. HAEC cultured on stiff hydrogels were stiffer (*p*<0.05) than those cultured on soft hydrogels (Suppl. Fig. 5A). In addition, integrin adhesion force was higher (*p*<0.05) in cells cultured on stiff vs. soft hydrogels (Suppl. Fig. 5B). Cells cultured on stiff hydrogels also exhibited increased heterogeneity, as reflected by HWHM values (Suppl. Fig. 5 A & B, table).

## Discussion

A major finding of this study is that pharmacological inhibition of LOX/LOXL enzymes ameliorates intimal stiffening induced by chronic exposure to d-flow. Of note, these effects were consistently observed in arteries from male and female mice. Thickening and stiffening of the intima in areas where the arterial tree undergoes geometric transitions (such as bifurcations, branch vessels, and curvatures) may constitute an initially adaptive response to changes in local hemodynamics. However, when combined with additional stressors such as metabolic imbalance, hypertension, or aging, altered blood flow patterns in these areas can lead to the formation of atherosclerotic plaque.^26, 27^ Results from this study suggest that reducing arterial wall stiffening per se could prevent the formation and rapid progression of atherosclerotic lesions. Therefore, identifying targets that are uniquely sensitive to arterial wall stiffening induced by persistent d-flow, in the absence of confounding atherogenic factors, could potentially advance the development new therapeutic approaches. These new strategies would complement existing cholesterol-lowering therapies for atherosclerosis.

Measurements performed just 2 days after PCL provided insights into the effects of acute exposure to d-flow in the absence of changes in the stiffness of the subendothelial matrix. Unexpectedly, the endothelium was softer in acutely ligated arteries of males. Acute exposure to oscillatory shear stress was shown to induce transient endothelial dysfunction, assessed by sustained stimulus flow-mediated dilation, in men but not in women.^28^ Studies *in vitro* have also identified sex-dependent responses of endothelial cells exposed to combinations of shear stress and substrate stiffness; overall endothelial cells from males were more sensitive than those from females to changes in the mechanical microenvironment.^29^ Further studies coupling analysis of mechanical and functional properties of the arterial wall will eventually elucidate the significance of sex-differences in the endothelial response to short-term disturbed blood flow *in vivo*.

Enzymes of the LOX/LOXL family play a key role in collagen and elastin crosslinking, thereby contributing to their mechanical properties, i.e., tensile strength and elasticity. The LOX/LOXL enzymes catalyze the oxidative deamination of lysine and hydroxylysine residues to generate highly reactive lysyl-aldehyde (allysine) which, in turn, undergoes spontaneous reactions with other lysine, hydroxylysine or allysine residues to create crosslinks.^30^ Previous research showed that a feed-forward loop involving extracellular matrix deposition and LOX/LOXL-dependent collagen crosslinking promotes atherosclerosis progression in hyperlipidemic, ApoE-deficient mice.^31^ In such context, arterial wall softening by the LOX/LOXL family inhibitor BAPN, attenuated atherosclerotic plaque formation.^31^ Here, we combined two weeks of PCL with continuous delivery of BAPN in wild type (C57BL/6J) mice fed regular chow. Our study, therefore, examined the direct contribution of LOX/LOXL activity to arterial wall stiffening induced by chronic d-flow without confounding effects of hypercholesterolemia or ApoE inactivation. In line with previous findings,^31^ the prevention of d-flow-induced intimal stiffening by BAPN administration suggests that extracellular matrix-targeted approaches that preserve or restore the mechanical homeostasis of the arterial wall may prove effective in reducing the burden of cardiovascular disease.

Alterations in the mechanical characteristics of the arterial wall can potentially influence the communication between endothelial cells and smooth muscle cells. For example, compared to the healthy endothelium of compliant arteries in young mice, the compromised nitric oxide release by the dysfunctional endothelium of stiff arteries in aged mice results in increased synthesis and secretion of LOX/LOXL enzymes by smooth muscle cells.^5^ These observations aligned with those of Jo and colleagues^11^ who, using the PCL model, demonstrated that persistent but not short-term exposure to d-flow impairs endothelium-dependent, nitric oxide-mediated arterial relaxation. The present observations, in conjunction with those of others,^11, 16^ suggest the existence of a mechanically driven feedback loop.

This loop entails the disruption of endothelial homeostasis by prolonged exposure to d-flow, which fosters stiffening of the arterial wall. This, in turn, exacerbates endothelial dysfunction. Significantly, links between endothelial stiffening and various indicators of endothelial dysfunction, including decreased release of nitric oxide,^32^ increased permeability,^33, 34^ elevated expression of cell adhesion molecules,^35^ and augmented leukocyte adhesion,^36^ have been well documented.

The present results suggest that BAPN administration blocks stiffening of the subendothelial stiffening induced by d-flow and restores, to a significant extent, the mechanical homeostasis of the endothelium. Nonetheless, we observed that the endothelium of ligated, BAPN-treated carotids was ∼2-fold stiffer than that of unligated, saline treated counterparts. Thus, in the absence of subendothelial matrix stiffening, chronic stress from d-flow may be sufficient to induce subtle changes in the mechanical phenotype of the endothelium. This finding is supported by recent work suggesting that endothelial stiffening may occur without changes in the stiffness of the arterial wall^37^ or the subendothelial matrix.^38, 39^ Indeed, Levitan and colleagues demonstrated that endothelial stiffening and dysfunction, arising from the combined effects of d-flow and dyslipidemia or aging, leads to early stages of atherogenesis.^38, 39^

While the deliberate inclusion of sex as a biological variable of importance in cardiovascular research has increased in recent years, much of the literature is built on studies excluding females or presenting results combining observations from males and females without rigorous testing for sex specific effects.^40, 41^ A strength of this study is the examination of males and females and inclusion of sex as a biological variable in data analysis. Our study shows comparable intimal stiffening in arteries from male and female mice exposed to persistent d-flow, an effect that was substantially mitigated by BAPN treatment in both sexes. Future studies examining distinctive alterations in molecular profiles and cellular phenotypes linked to persistent d-flow, coupled with strategies aimed at preserving or restoring the mechanical homeostasis of the arterial wall, hold the potential to unveil novel therapeutic avenues for atherosclerotic cardiovascular disease.

## Limitations of the study

One constraint within this study relates to the absence of an experimental group subjected to acute d-flow in conjunction with BAPN treatment. Technical limitations hindered the acquisition of force curves from unligated RCA+BAPN carotid samples displaying a softer-than-anticipated nature. Further investigations might be needed to comprehensively understand the immediate and prolonged impacts of BAPN treatment on the mechanical properties of the intima.

The present investigation underscores the potential of inhibiting LOX/LOXL enzymes as an effective strategy to offset intimal stiffening. Subsequent inquiries that specifically address the contribution of LOX/LOXL enzymes to chronic d-flow exposure could pave the way for innovative, precision-targeted therapies against arterial stiffening. Lastly, the potential influence of inherent stiffening in smooth muscle cells, an aspect not explored within this study, could provide supplementary insights into the mechanisms underlying arterial wall stiffening stemming from persistent d-flow exposure.

## Conclusions

This study was primarily designed to test the hypothesis that inhibition of arterial wall stiffening neutralizes the deleterious effects of chronic d-flow on the mechanical properties of the arterial intima. Whereas acute d-flow appears to exert minimal effects on the mechanical properties of the arterial intima, persistent d-flow leads to remarkable stiffening of both the endothelium and the subendothelial matrix. Intimal stiffening induced by persistent d-flow can be averted through pharmacological inhibition of the extracellular matrix-modifying enzymes of the LOX/LOXL family.

